# A TOPOLOGICAL/ECOLOGICAL APPROACH TO PERCEPTION

**DOI:** 10.1101/086827

**Authors:** James F. Peters, Arturo Tozzi, Sheela Ramanna

## Abstract

During the exploration of the surrounding environment, the brain links together external inputs, giving rise to perception of a persisting object. During imaginative processes, the same object can be recalled in mind even if it is out of sight. In this proof of concept study, Borsuk’s theory of shape and the Borsuk-Ulam theorem provide a mathematical foundation for Gibson’s notion of persistence perception. Gibson’s ecological theory of perception accounts for our knowledge of world objects by borrowing a concept of invariance in topology. A series of transformations can be gradually applied to a pattern, in particular to the shape of an object, without affecting its invariant properties, such as boundedness of parts of a visual scene. High-level representations of objects in our environment are mapped to simplified views (our interpretations) of the objects, in order to construct a symbolic representation of the environment. The representations can be projected continuously to an ecological object that we have seen and continue to see, thanks to the mapping from shapes in our memory to shapes in Euclidean space.

---

Theoretical physics teaches us that the intimate micro-structure of the world consists of elements in perpetual movement, interacting with each other in a framework of probabilities, energy fields and vacuum. However, when we see a segment in a visual scene in the environment, we perceive elements seemingly melted together in a single complex of sensations. To make an example, we are able to detect, in the indistinctness of a rural scene at sunrise time, an increasingly distinct world of trees, hills, valleys and moving particles, e.g., birds flying from one tree to another. In effect, we appear to be sewing pieces of a changing scene together. How does the brain join different significant elements, giving rise to a single, stable perception of a scene? How does the brain imagine or recall objects that are out of sight? These two problems need to be tackled not at the microscopic atomic or sub-atomic level, nor at the galactic level, but at an intermediate macroscopic level where living beings stand. A noteworthy approach to understanding the mechanisms of direct perception is to start at the ecological level. Indeed, this article introduces a shape-based explanation of J.J. Gibson’s theory of persistence perception (Gibson, 1950). Throughout the second half of the twentieth century, James J. Gibson (1966, 1971, 1979) developed a unique theory in perceptual science, namely the ecological theory of perception (ETP), which stresses the importance of the relationships of the individual and the environment (Heft, 1997). The foundation for perception is ambient, ecologically available, direct information, as opposed to peripheral or internal sensations. The human (and/or animal) individual is embedded in the surrounding environment and the perception is strictly linked to searching movements, because changing awareness of the connectedness of the components of a scene occurs *in situ*, and not in a static context. The environment contains real, perceivable opportunities (called “affordances”) for an individual’s actions. Affordances are properties of the perceived environment (Gibson, 1979; Gibson, 1986; Lu, 2013) that provide paths to merged structures, shapes and actions. For example, the more chances children are given to directly perceive and interact with their environment, the more affordances they discover, and the more accurate their perceptions become over the passage of time. Perceiving is essentially exploratory (Heft, 1997). It consists in the acquisition of objective knowledge, by assembling diverse views of an object to form its abstract representation. In effect, a perceptual system is formed by vision, movements of the eyes, head and entire body. Gibson argued that perception of persisting surfaces depends on the perception of specific structures, invariant over time. Recognition (perceiving) objects entails continuous mappings between our representations of objects and in-the-world ones. In effect, perception can be explained by our discovery of equivalent configurations (form, shapes) of objects (Heft 1997; Rock, 1983).

Visual information is carried in two separate pathways in our brain (Mishkin et al., 1983). Visual identification of objects is enabled by the multisynaptic corticocortical ventral pathway, which interconnects the striate, prestriate, and inferior temporal areas. On the other side, visual location of objects is made possible by the dorsal pathway, which runs dorsally, interconnecting the striate, prestriate, and inferior parietal areas. Therefore, the visual system separates processing of an object’s form and color (“what”) from its spatial location (“where”) (Rao et al., 1997). Recent findings suggest that nervous structures process information through topological as well as spatial mechanisms. For example, it is has been hypothesized that hippocampal place cells create topological templates to represent spatial information (Dabaghian, 2014; Arai, 1014; Chen, 2014). Developments in studies which consider the role of embodied cognition and action in psychology can be seen to support this basic position about hippocampal place cells. In particular, in this article, we show that novel incarnations of the “classical” Borsuk-Ulam theorem lead to a better comprehension and assessment of several features of direct perception and imagination.

## MATERIALS AND METHODS

### Borsuk-Ulam theorem

Topology, which assesses the properties that are preserved through deformations, stretchings and twistings of objects, is a underrated methodological approach with countless possible applications (Willard, 1970; Krantz, 2009; Manetti, 2015). The Borsuk-Ulam theorem or BUT, cast in a quantitative fashion which has the potential of being operationalized, is a universal principle underlying a number of natural phenomena. In particular, we will show that BUT and its variants allow the assessment of perceptual features in terms of affinities and projections from real spaces to representations in feature spaces. It is noteworthy that BUT and its variants talk about projections, connections, not about cause-and-effect relationships (Tozzi and Peters, 2016a). Briefly, BUT states that, if a single point on a circumference projects to a higher spatial dimension, it gives rise to two antipodal points with matching description on a sphere, and vice versa (Figure 1A) (Borsuk, 1933; Marsaglia, 1972; Matoušek, 2003; Beyer, 2004). This means that the two antipodal points are assessed at one level of observation in terms of description, while a single point is assessed at a lower level (Tozzi 2016b), i.e., point location vs. point description. Points on a sphere are “antipodal”, provided they are diametrically opposite (Henderson, 1996). Examples of antipodal points are the poles of a sphere. This means, e.g., that there exist on the earth surface at least two antipodal points with the same temperature and pressure. BUT looks like a translucent glass sphere between a light source and our eyes: we watch two lights on the sphere surface instead of one. But the two lights are not just images, they are also real with observable properties, such as intensity and diameter.

### Variants of the Borsuk-Ulam theorem

The concept of antipodal points can be generalized to countless types of system signals. The two opposite points can be used not just for the description of simple topological points, but also of lines, or perimeters, areas (Figure 1B), regions, spatial patterns, images, temporal patterns, movements, paths, particle trajectories (Figure 1C) (Borsuk, 1969; Peters and Tozzi, 2016b), vectors (Figure 1D), tensors, functions (Figure 1E), algorithms, parameters, groups, range of data, symbols, signs, thermodynamic parameters (Figure 1F), or, in general, signals. If we simply evaluate nervous activity instead of “signals”, BUT leads naturally to the possibility of a region-based, not simply point-based, brain geometry, with many applications. A region can have indeed features such as area, diameter, illumination, average signal value, and so on. We are thus allowed to describe system features as antipodal points on a *s*phere (Weeks, 2004; Peters, 2016). Descriptively similar points and regions do not need necessarily to be opposite, or on the same structure (Figures 2A and 2B (Peters, 2016; Peters and Tozzi 2016a). Therefore, the applications of BUT can be generalized also on non-antipodal neighbouring points and regions on an sphere. In effect, it is possible to evaluate matching signals, even if they are not “opposite”, but “near” each other. As a result, the antipodal points restriction from the “standard” BUT is no longer needed, and we can also consider regions that are either adjacent or far apart. And this BUT variant applies, provided there are a pair of regions on the sphere with the same feature value. We are thus allowed to say that the two points (or regions, or whatsoever) do not need necessarily to be antipodal, in order to be labeled together (Peters 2016). The original formulation of BUT describes the presence of antipodal points on spatial manifolds in every dimension, provided the manifold is a convex, positive-curvature structure (*i.e*, a ball). However, many brain functions occur on manifolds endowed with other types of geometry: for example, the hyperbolic one encloses complete Riemannian *n*-manifolds of constant sectional curvature -1 and concave shape (*i.e*, a saddle) (Sengupta et al., 2016). We are thus allowed to look for antipodal points also on structures equipped with kinds of curvature other than the convex one (Figure 2C) (Mitroi-Symeonidis, 2015). Or, in other words, whether a system structure displays a concave, convex or flat appearance, does not matter: we may always find the points with matching description predicted by BUT (Tozzi, 2016). A single description on a plane can be projected to higher dimensional donut-like structures (Figure 2D), in order that a torus stands for the most general structure which permits the description of matching points. For further details, see Peters and Tozzi (2016b).

**Figure 1.**
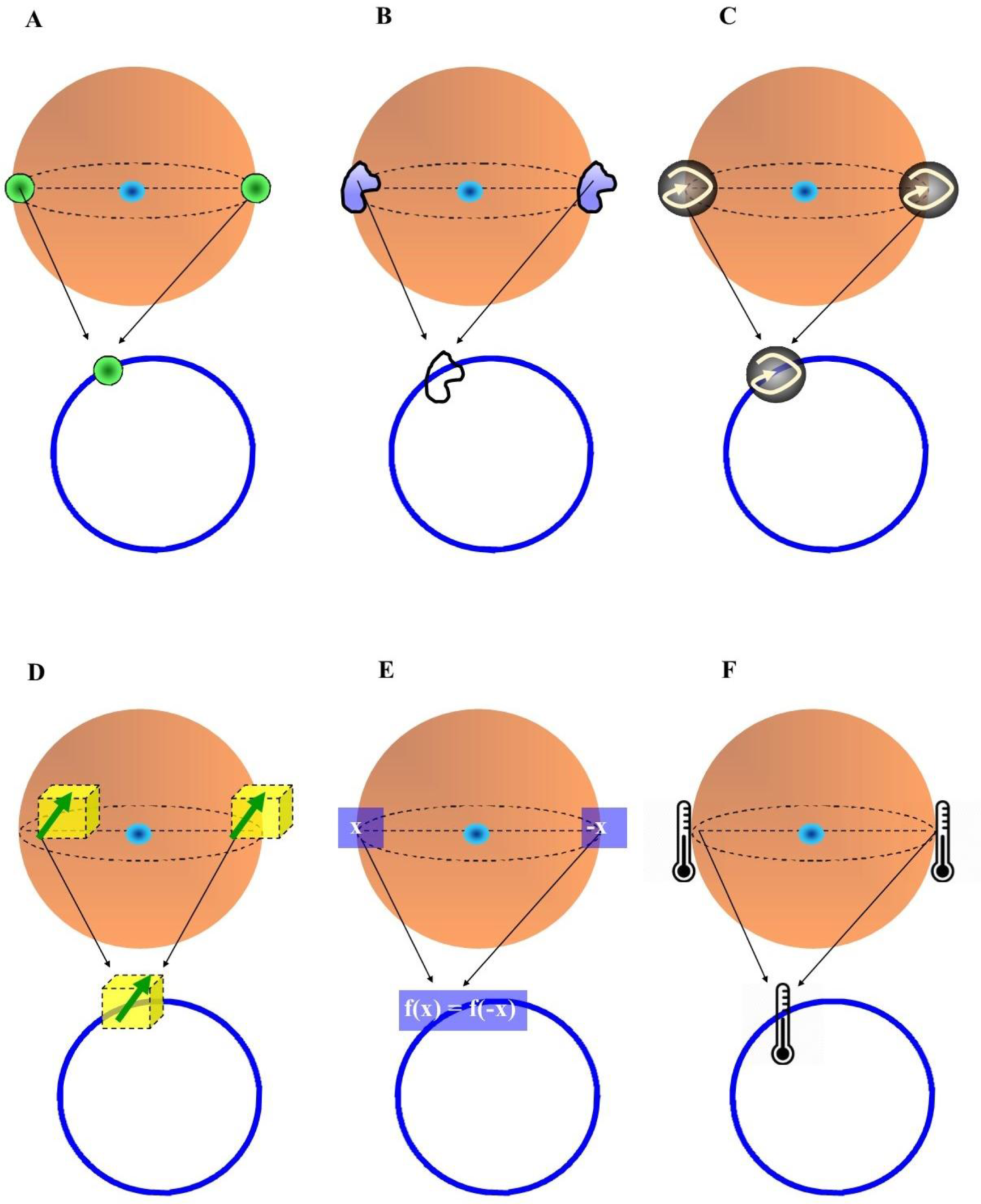
Possible types of antipodal features in Borsuk Ulam theorem (BUT). The original formulation of the BUT, with two antipodal points, is illustrated in Figure 1A. **Figures 1B-1F** display, respectively, two antipodal shapes, trajectories, vectors, functions and temperatures. Each set of antipodal features on a sphere stands for a single feature on a circumference, if we evaluate it just one dimension lower.

**Figur 2.**
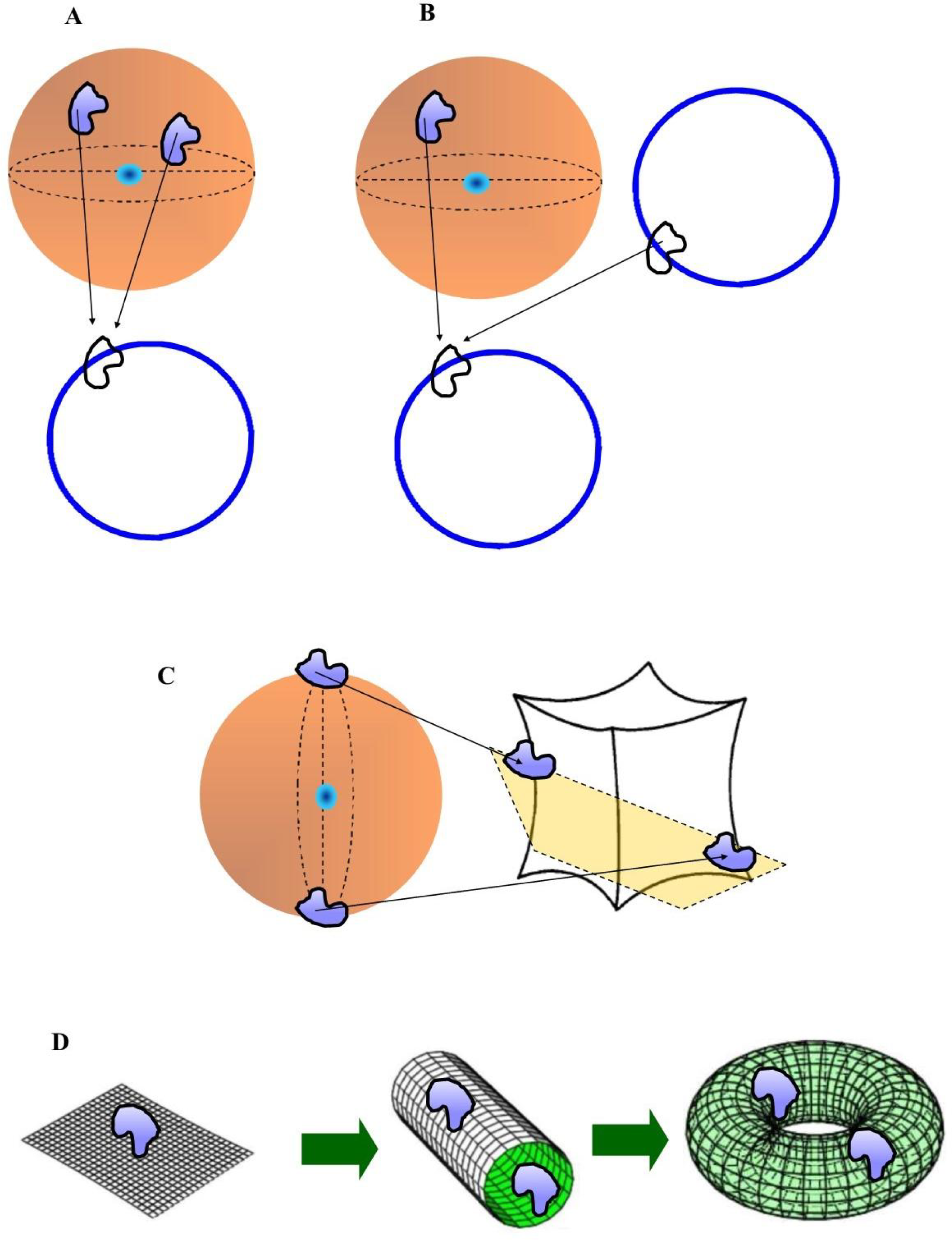
Two features with signal matching do not need necessarily to be antipodal. Indeed, the applications of the BUT can be generalized also on non-antipodal points on the same sphere (Figure 2A), of non-antipodal points lying on two different structures (Figure 2B). Figures 2C and **2D**: BUT applies for structures with different types of curvatures. See the main text for further details.

Although BUT has been originally described just in case of *n* being a natural number which expresses a structure embedded in a spatial dimension, nevertheless the value of *n* in the brain sphere can also stand for other types of numbers. The BUT can be used not just for the description of “spatial” dimensions equipped with natural numbers, but also of antipodal regions on brain spheres equipped with other kinds of dimensions, such as a temporal or a fractal one. This means, e.g., that spherical structures can be made not just of space, but also of time. The dimension *n* might stand not just for a natural number, but also for an integer, a rational, an irrational or an imaginary one. For example, *n* may stand for a fractal dimension, which is generally expressed by a rational number. This makes it possible for us to use the *n* parameter as a versatile tool for the description of systems’ features.

Furthermore, matching points (or regions) might project to lower dimensions on the same structure (Figure 3A). A sphere may map on itself: the projection of two antipodal points to a single point into a dimension lower can be internal to the same sphere. In this case, matching descriptions are assessed at one dimension of observation, while single descriptions at a lower one, and vice versa. Such correlations are based on mappings, e.g., projections from one dimension to another. In many applications (for example, in fractal systems), we do not need the Euclidean space (the ball) at all: a system may display an intrinsic, *internal* point of view, and does not need to lie in any dimensional space (Weeks, 2002). Therefore, we may think that the system just does exist by - and on – itself.

**Figure 3.**
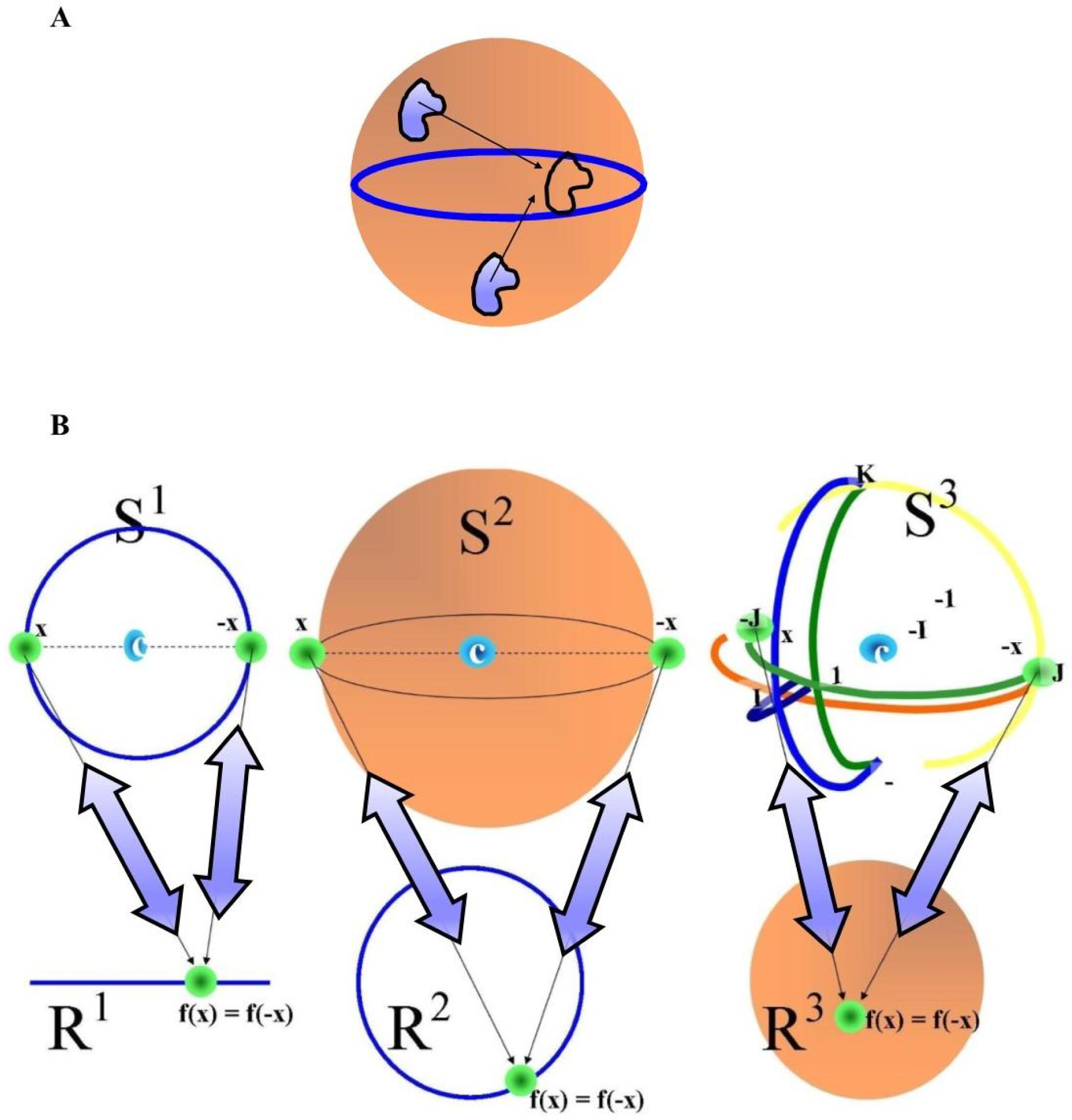
shows how a sphere may map on itself. See the main text for further details. Figure 3B displays the BUT acting on structures of different dimensions: a 2-D circle, a 3-D sphere and a 4-D hypersphere. A symmetry, e.g., two antipodal points in higher dimensions, is said to be “broken” when it maps to a single point in one dimension lower. Note that the mapping is reversible from higher to lower dimensions, and vice versa (blue double-sided arrows).

### The foremost role of symmetries

Symmetries are real invariances that underline the world and occur at every level of systems’ organization (Weyl, 1982). Bilinear forms of geometrical structures manifest symmetrical properties (Weyl, 1918). In other words, affine structures preserve parallel relationships. Symmetries are the most general features of mathematical, physical and biological entities and provide a very broad approach, explaining also how network communities integrate or segregate information. It must be emphasized that symmetries are widespread and may be regarded as the most general feature of systems, perhaps more general than free-energy and entropy constraints too. Indeed, recent data suggest that thermodynamic requirements have close relationships with symmetries. The interesting observation that entropy production is strictly correlated with symmetry breaking in quasistatic processes paves the way to use system invariances for the estimation of both the free energy of metastable states, and the energy requirements of computations and information processing (Roldán, 2014). Thus, giving insights into symmetries provides a very general approach to every kind of systems function. A symmetry break occurs when the symmetry is present at one level of observation, but “hidden” at another level. A symmetry break is detectable at a lower dimension of observation (Figure 3B). Thus, we can state that single descriptions are broken (or hidden) symmetries, while matching descriptions are restored symmetries. In other words, a symmetry can be hidden at the lower dimension and restored when going one dimension higher. If we assess just single descriptions, we cannot see their matching descriptions: when we evaluate instead systems one dimension higher, we are able to see their hidden symmetries. In sum, symmetries, single descriptions and matching descriptions are the common language among different real sciences.

In sum, BUT displays versatile formulations that can be modified in different guises, in order to achieve a wide range of uses in neuroscience, such as multisensory integration, neural histological images tessellation, brain symmetries, Kullback-Leibler divergence during perception, detection of a brain functional 3-sphere. For further details, see Tozzi (2016a and 2016b). In the next paragraphs, we will focus on the role of BUT and its variants in direct perception and object recalling, taking into account Gibson’s incisive observations.

### Perception of shapes and perception as shape mapping

In such a theoretical context of perception, the Borsuk-Ulam theorem helps elucidate the tenets of the ETP and explains how we see an object and how we imagine it (Tozzi and Peters, 2016b). Briefly, BUT explains how high-level representations of objects in our environment are typically mapped to simplified views (our interpretations and coalescences) of the objects. And these mappings described by BUT can sometimes be reversed (*inversed*) to achieve a form of pullback from descriptions to sources of descriptions, from a simplified view to multiple views of the same object. Indeed, a new form of shape theory (called homotopy) discovered by K. Borsuk makes it possible to assess the properties that are preserved through deformation, stretching and twisting of objects (Beyer, 2004; Manetti, 2015). Homotopy is a theory of shape deformation (Borsuk, 1971; Borsuk and Dydak, 1980), e.g., how some shapes can be deformed into other shapes. In this context, the term “shape” means “exterior form” and a “deformation” is a mapping from shape into another one. A classical example is the deformation of a coffee cup into a torus. The combination of various forms of BUT and homotopy theory provides a methodological approach with countless possible applications, especially in helping us understand perception and how we acquire visual imagination. The theory of shape, in simple terms, focuses on the global properties of geometric objects such as polyhedra and tori, neglecting the complications of the local structures of the objects (Borsuk and Dydak, 1980). What shape theory and BUT tell us about visual image acquisition echoes the Gibsonian view of perception: *cognitive processes relating to perception, storage, retrieval and reorganization interact with memory structures and construct a symbolic representation of the environment*. For example, mesh view in **Figure 4** can be viewed as visualization of one among many symbolic views of Leonardo Da Vinci’s Mona Lisa painting. In effect, an individual perceives [constructs] the features and events of the environment specified by this [pickup] information from the environment (Heft, 1997). In particular, BUT variants and Borsuk’s theory of shapes are closely allied to Gibson’s view of *persistence of perception* (Gibson, 1979; Natoulas, 1992). That is, our perception of an object continues, even when the object is out of sight. This can concisely explained by viewing regions on the surface of a hypersphere as multiple representations of object shapes, mapped continuously to an ecological object that have seen and continue to see. It occurs thanks to the continuous mapping from shapes in our memory to shapes in Euclidean space. In effect, persistence perception can be viewed as signals matching, real-scene visual signals that are collectively the umbra of physical shapes. To make an example, **Figure 4** displays the Mona Lisa painting impinging on the optic nerves: we map it to similar shape representations, such as the Mona Lisa mesh. Gibson accounted for our knowledge of world objects by borrowing a concept of invariance in topology: a series of transformations can be endlessly and gradually applied to a pattern without affecting its invariant properties (Gibson, 1950). And, principal among the properties of world objects, is shape and the acquisition of persistent perception of object shapes, which we nicely explain with BUT variants.

**Figure 4.**
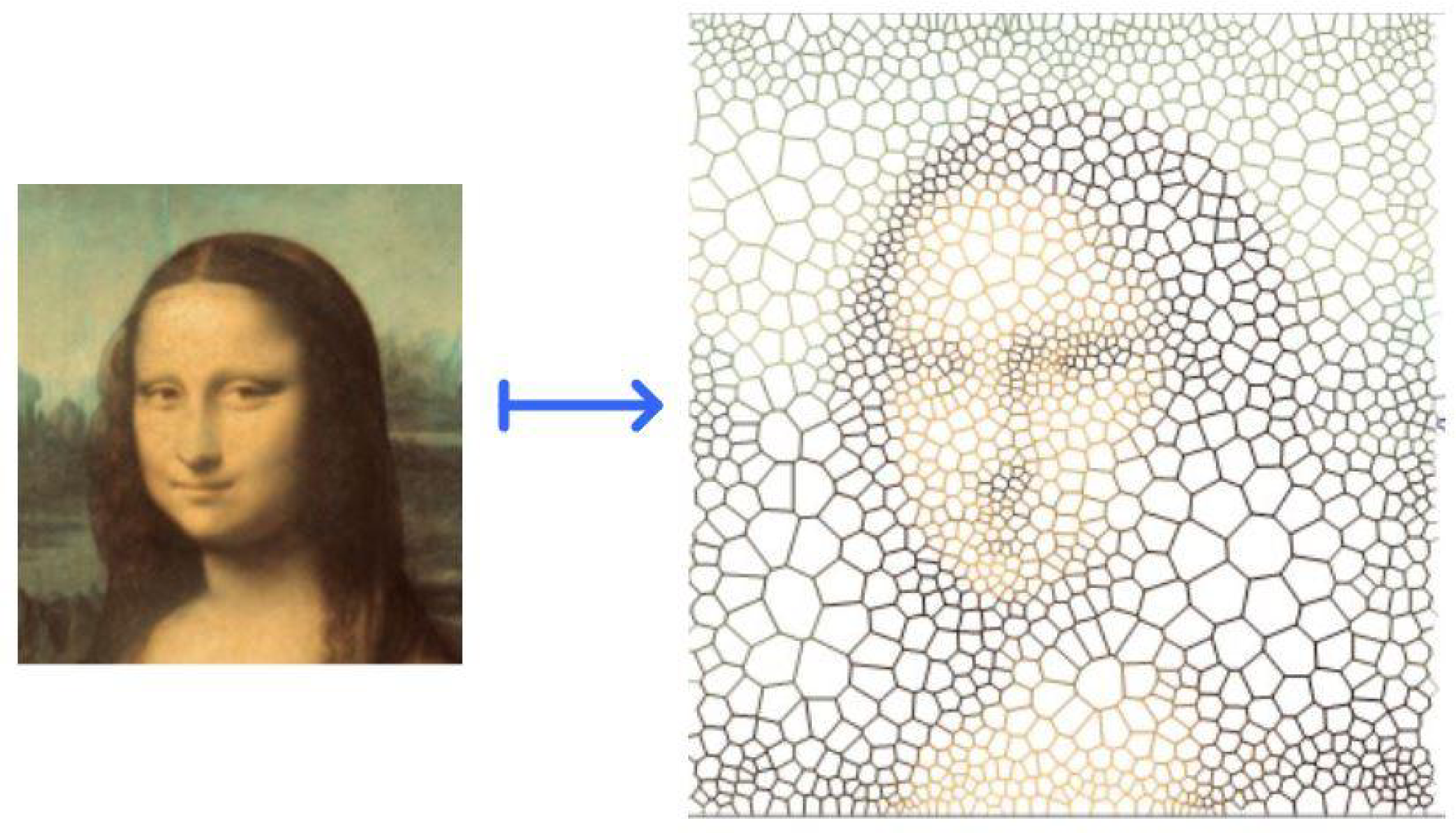
Mona Lisa shape deformably mapped to Mona Lisa Exterior Mesh Shapes. This mapping illustrates nascent Gibsonian persistence perception: we abstract away from the very complex structures in Da Vinci’s painting to arrive a simpler, geometric view of the image as a collection of familiar convex polygons.

### A topological approach to Gibson’s “ask not what’s inside your head, but what your head’s inside of”

We start from the simplest and most natural standpoint: an individual (either human or animal) embedded in his environment (Russo-Krauss, 2015). The environment stands for what surrounds the organisms that perceive and behave, in order that both are joined together in a inseparable couple. In topological terms, we may say that the individual is embedded in a spatial three-dimensional ball. Thus, the individual stands in front of the surrounding spatial environment, shaped in guise of a sphere. The environment is not the one of the “traditional” physics: indeed, it can be described through – and is made of - the triad of “medium”, “substance” and “surfaces” perceived by the individual. The ecological surface stands for the “classical” geometric plane, while the ecological medium for the standard geometrical space. In such a guise, the object features that the individual perceives are just the interface between surfaces, in order that information is embedded into the light (natural or artificial) surrounding them. The most stable, long-lasting perception of the individual is the base upon which he stands, e.g., the ground, the horizontal surface under his feets. When the individual raises his eyes, he perceives the sky, or the ceiling. The middle zone between the ground and the sky is the medium. The objects, e.g., something which can be handled, stand vertically in the medium. The permanent objects are just surface features which can be perceived on long timescales. For our purposes, it is noteworthy that the human (and also animal) individual splits the environment, in which he moves and finds novel affordances, in three components: the ground and the sky, which join together at the horizon, and the intermediate zone, the medium. We are thus allowed to split the three-dimensional ball into three slices: the sky on the top, the ground on the bottom and the intermediate zone in the middle. In topological terms, the brain splits the environment in different closed subsets. In this framework, a theorem, closely linked to the Borsuk-Ulam theorem, comes into play: the Lusternik–Schnirelmann theorem (LST). It states that, if a sphere is covered by n+1 open sets, then one of them contains a pair of antipodal points. In other words, every time you split a sphere in three parts, there is always one of them which contains an entire diameter, where the antipodal points lie (Figure 5A). With this theorem, we are guaranteed at least a pair of exactly opposite points on a sphere (Dodson and Parker, 1997). According to LST, one of the three slices must necessarily contain some antipodal points, e.g., surface features with matching description. When an individual moves around and looks for something (i.e., an object), his brain focuses on one of the three slices of the n-sphere. If we state that the individual attention is driven by one of the slices, we may term such slice as the one containing antipodal points. In other words, the individual needs to find, in one of the three slices in which he separates his environment (the sky, the ground and the medium), a pair of antipodal points equipped with matching description. The latter, by an ecological point of view, share something perceivable in common. Note that, according to the forms of the BUT, the points do not need necessarily to be antipodal. It means that two points embedded in one of the three closed sets do not need to be opposite to be described together. The antipodal points, according to one of the BUT variants, do not need to be simple *points*, but they could have different features. They could be spatial objects or shapes, but also matching movements, or they may display a temporal scale. In the latter case, the shapes are not called objects, but events.

**Figure 5A.**
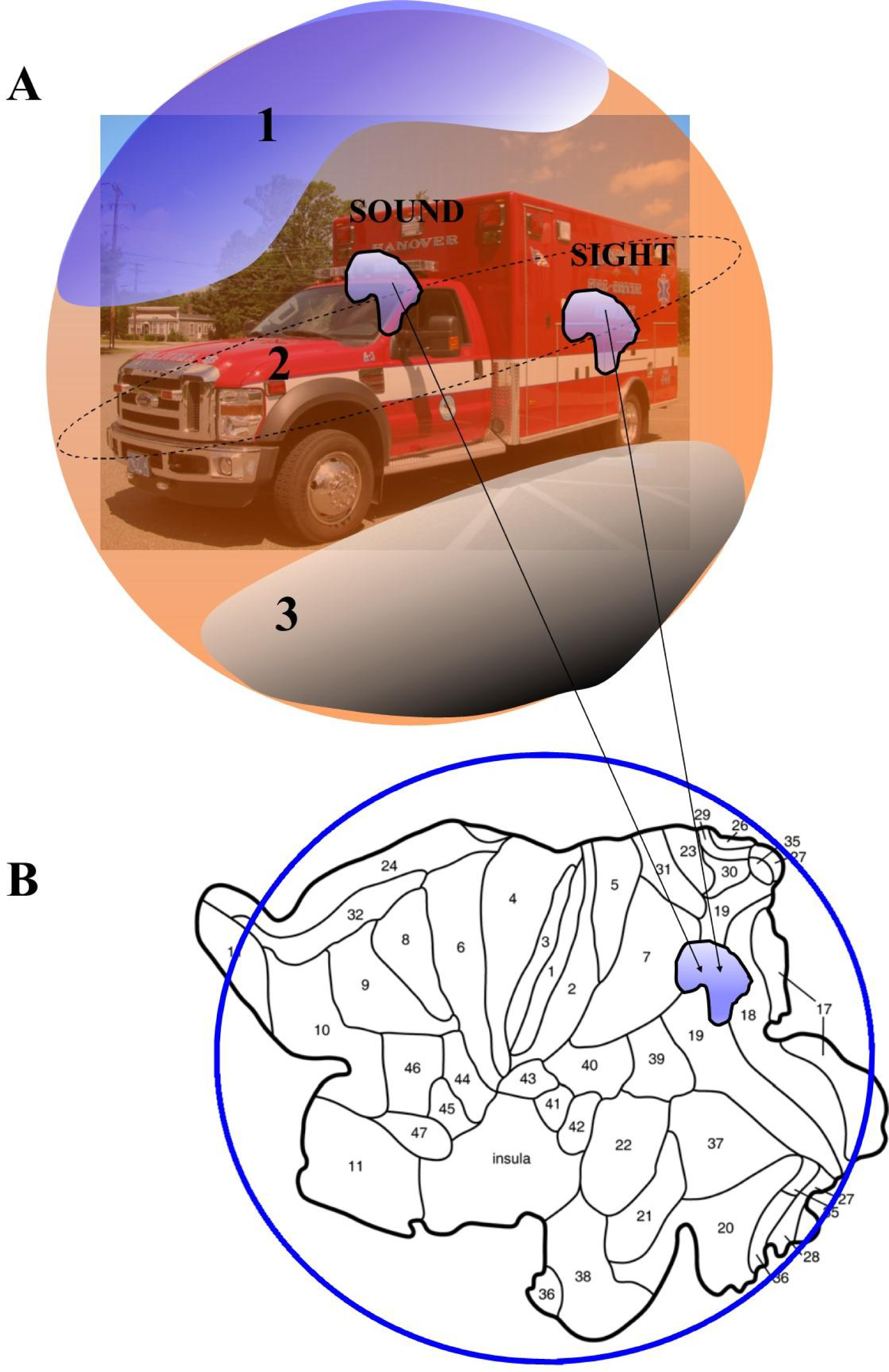
The three closed regions predicted by the Lusternik–Schnirelmann theorem are named 1, 2 and 3. Once achieved two points with matching description embedded into one of the three closed regions (in this case, the region 2), they can be projected (Figure 5B) to a single point onto a 2D brain model (modified from Van Essen, 2005).

## RESULTS

Once we have established a topological framework for objects in the environment, how does the process of perception occur in our brain (mind)? When two antipodal points (or functions) are joined together according to BUT or its variants, they are perceived together, in order that the individual sees (or hears, or touches) a complex of sensations representing the object. According to one of the BUT features (the mapping from an higher to a lower dimension), the dimensions of the surrounding environment need to be reduced when they “enter” our brain. To make an example, we will take into account the multisensory integration, a mental phenomenon occurring when two stimuli from different cues are perceived together (e.g., the view of a running ambulance and the sound of its siren, Figure 5A). In a topological view, BUT (and its variants) nicely applies to multisensory integration: two environmental stimuli from different sensory modalities display similar features, when mapped into cortical neurons. To make an example, an observer stands in front of the surrounding environment. An ambulance is embedded in the environment. The observer perceives, through his different sense organs such as ears and eyes, the sounds and the movements of the ambulance. The latter stands for an object embedded in the three-dimensional sphere. The two different sensory modalities (sounds and movements) stand for non-antipodal points on the sphere’s circumference. Even if objects belonging to opposite regions can either be different or similar, depending on their features (Peters, 2016), however they must share the same features. In our case, both sounds and movements come from the same object embedded in the sphere, i.e., the ambulance. The two non-antipodal points project to a two-dimensional layer, the brain cortex - where multisensory neurons lie - and converge into a single multisensory signal (Figure 5B). According to the dictates of the BUT, such a single point contains a “melted” messages from the two modalities, which takes into account the features of both (Tozzi and Peters, 2016b).

## CONCLUSIONS

Our paper provides a link between the Ecological Theory of Perception (Gibson ETP) and the topological framework of BUT and its variants, able to explain the mechanisms of perception. Although Gibson stated that space is a myth, a ghost, a fake produced by geometry, a topological approach sheds new light on his own theory. According to this novel conception, the individual’s brain (or mind) is forced to coalesce together some components of the environment, in a complex interaction between external affordances and the motivated humans who perceive them. The melting of parts of the environment into a single perception is thus compelled, and is not a free choice made by the individual. The brain needs to perceive different elements together and cannot split them, because the perception may occur and operate just in this way. Contrary to the standard theories of perception, which emphasize the input processing and an helmotzian distinction between fixed, innate sensations and variable, learned perceptions, ETP proposes a novel approach. A mutual relationship occurs between the individual and the environment. The perceptive experience is direct and immediate. Informations are directly taken from the environment, with no mediations. According to ETP, the real word is the phenomenic one. The ETP concept of a smart mechanism able to extract the invariants from the input stream means that the active subject puts in relationships and connexions the objects through his senses, with no need for mental processing in order to build them. The truth corresponds to the immediate perception, which is a *datum* in our brain. Topology nicely explains this interesting feature which is common not just to perceptions, but also to higher mental activities. Indeed, the antipodal point do not stand just for spatial features, such as an object, but also for temporal sequences. It means that the brain is constrained to link together not just spatial elements in an object, but also temporal events in a concept, an idea, a proposition, a hope, a desire and so on. Furthermore, the individual can be embedded not just in a spatial n-sphere, but also in a n-sphere in which n stands for the time, according to the dictates of the BUT variants. Also many spheres with different times are possible: it means that the individual is “constrained” to link past and future events, it is not a choice of his own.

The evolutionary advantage is self-evident: the perception of different elements is useless by itself, while the perception of a complete object, or of a concept or an idea, is mandatory in order to survive in an explorable environment, full of possibilities, but also of dangers. In sum, the need to join things together in a single perception is mandatory for our brain (mind). It is important to emphasize that the antipodal points with matching description do not need to be causally correlated: their relationship is a topological one, meaning that the surface features of an object are “linked” together in a single complex of sensations by projections, affine connexions and proximity. In other words, the concept of *connexion* means that the joined parts if the environment are not necessarily in causal relationship, rather they are simply functionally correlated. The individual, with the habituation, learns, from childhood forward, to join together the elements which are more useful for his surviving.

“Perception is based on information, not on sensations” (Gibson, 1979). This means that ETP, rather than sensation-based, is information-based, because emphasizes an analysis of the environment (in terms of affordances), and the concomitant specificational information that the organism detects about such affordances. The human behaviour is radically situated. In other words, you cannot make predictions about human behaviour unless you know what situation or context or environment the human in question was in. Individuals stand in an ecological relation to the environment, such that to adequately explain some behaviour it is necessary to study the environment or niche in which it took place and, especially, the information that “epistemically connects” the organism to the environment. Thus, an appropriate analysis of the environment, made in terms of BUT and its variants, becomes crucial for an explanation of perceptually guided behaviour. Gibson’s work strengthens and brings to the front the primary question of “what” is perceived, before questions of mechanisms and material implementation are introduced (Rao et al., 1997). Together with a contemporary emphasis on dynamical systems theory as a necessary methodology for investigating the structure of ecological information, the Gibsonian approach, particulary in the light of the modern tools of algebraic topology, has maintained its relevance and applicability to neuroscience.

## Acknowledgements

The authors would like to thank Rosa Del Prete for her precious suggestions.

